# EGL-19 and EXP-2 Ion Channels Modelling within C. Elegans Pharyngeal Muscle Cell Allows Reproduction of CA2+ Driven Action Potentials, Both Single Events and Trains

**DOI:** 10.1101/228650

**Authors:** Andrey Yu. Palyanov, Khristina V. Samoilova, Natalia V. Palyanova

## Abstract

One of the current problems at the interface between neuroscience, biophysics, and computational modeling is the reverse-engineering and reproduction of *Caenorhabditis elegans* using computer simulation. The aim of our research was to develop the computational models and techniques for solving this problem while participating in the international open science OpenWorm Project. We have suggested models of a typical *C. elegans* neuron and a pharyngeal muscle cell, which were constructed and optimized using the NEURON simulation environment. The available experimental data about EGL-19 and EXP-2 ion channels allowed the model of a muscle to reproduce the action potential time profile correctly. Also, the model of a neuron reproduces quite accurately the mechanism of neural signal transmission based on passive propagation. We believe our models to be promising for better representing the specifics of various nervous and muscular cell classes when adding the corresponding ion channel models. Moreover, they can be used to construct the networks of such elements.

## 1. Introduction

The idea of reverse-engineering of the brain and its reproduction as computer simulation attracts scientists since the early days of computers. It is not surprising that the human brain comprising 98 billion neurons is still out of reach, although Human Brain Project^a^ and Human Connectome Project^b^ have been started recently. How much time and resources will it take, and will the project succeed? It seems highly reasonable to test the similar methodology and techniques on the most simple and well-studied organism to investigate whether the model works or not.

The only animal with almost completely determined *connectome* (a comprehensive map of neural connections at the level of individual neurons, synapses, gap junctions, connections to muscle and sensory cells etc) is a tiny free-living soil nematode *C. elegans* with an extremely small nervous system. The pioneering investigation of *C. elegans* in terms of neurobiology by J.G. White and colleagues^1^ revealed its relatively simple anatomy: the body is composed of 959 cells, 302 of which are neurons and 95 are body wall muscles. In spite of its anatomical simplicity, it displays surprisingly complex behavior: it was shown that *C. elegans* possesses the behavioral plasticity, learning and memory^2,3^.

Many laboratories all over the world work on *C. elegans* nervous system investigation and computational modelling, including the OpenWorm project^c^ aimed at creating the *in silico* C. elegans nematode^4^. Consideration of such a system might be really helpful for building the reliable background for further research and development. In this work we focused on the pharynx of *C. elegans* – a muscular pump which rhythmically contracts to grind and transport bacteria so it can be further digested in the intestine. Pharyngeal neuro-muscular system is almost isolated from the rest nervous system, it can function autonomously and contains significantly less elements than the whole neuro-muscular system, just 20 muscle cells and 20 neurons^d^.

A neuron usually consists of a cell body and thin long processes – single axon and commonly multiple dendrites. Neurons interact with each other via electrochemical signals which can be transmitted between them through synapses and gap junctions. Usually in higher multicellular organisms nervous signals travel along neurites via action potentials (APs) without loss of amplitude. This mechanism is fast, active and consumes energy. Another type of propagation is called electrotonic conduction. Such signals propagate slowly and fade as they spread along neurites. Usually they travel for distances no longer than a few millimeters. However, their amplitude at neuron endings is still high enough to trigger the release of a neurotransmitter and activate a post-synaptic neuron^5^.

An important role in regulation of neural, muscular or any other living cell membrane potential belongs to ion channels. There is a huge variety of them: ion-specific, voltage-gated, mechanosensitive and many other types are known. For example, voltage-gated Na+ and *K*+ define the shape and properties of recently mentioned APs. The nervous system of *C. elegans* lacks classic APs due to the absence of fast voltage-gated Na+ channels. They are absent in the genome of *C. elegans* and not detected via the electrophysiologic studies. It is thus supposed that the functions which are carried out in the nervous systems of other organisms by means of APs are implemented in *C. elegans* through gradual depolarization and electrotonic conduction. Its nervous system and ion channels (quite a common set, with the exception of Na+ voltage-gated one) are presumably adapted to this unusual mode of functioning^6^. *C. elegans* muscle cells are capable of generating APs, although quite unusual – long-lasting and driven by Ca^2+^ instead of Na+. A qualitative description of the major ion channels (CCA-1, EGL-19 and EXP-2) responsible for the stages of the pharyngeal AP is available^7^, as well as electrophysiologic recordings of APs.

Our aim is to reproduce the major mechanisms, including ion channels, neuro-physiologic parameters and morphology, which are necessary for a basic neurobiologically reasonable simulation which can be further enhanced with more sophisticated mechanisms. In this paper we address the following issues:

- Collection of available experimental data about electrophysiologic properties and morphology of a typical neuron and a pharyngeal muscle cell.
- Search for appropriate existing models of related ion channels to use them as entry points for modification and optimization.
- Development of a general neurobiologically grounded model of a neuron and of a pharyngeal muscle cell – the basis for further construction of specific cell types and networks composed of these elements.
- Nearly perfect quantitative reproduction of the shape of a pharyngeal muscle cell AP – obtained for the first time for this type of cells.

## 2. Methods and algorithms

In the field of simulation of biological neural networks there are two acknowledged leaders, the NEURON^e^ and the GENESIS^f^, with nearly equivalent capabilities. Our choice of the NEURON simulation environment^8^ was determined by a fact that it provides a Python interface to its functions and data structures. In our case it is necessary for implementation of data exchange between the model of neural network and the model of *C. elegans* physical body and habitat (which we developed recently^9^). Also NEURON includes a built-in programming language HOC and an ability for construction of custom models of ion channels and other cellular mechanisms via Neuron Model Description Language (NMODL) expanding NEURON’s standard repertoire. All the source codes for the channel models, cell models and for their optimization, were written in HOC and NMODL and are available on request.

### 2.1. On electrophysiology of C. elegans neurons

Recently we have collected and analyzed morphological data about *C. elegans* neurons and electrophysiologic parameters^10^. Typical *C. elegans* neuron is very small – the size of soma being about 2–3 *μm* and neurites usually about 0.1–0.5 *μ_m_* in diameter and up to 1 mm long. Its specific membrane resistance *R_m_* (resistance times unit area) was reported to be within the range of 61… 251 kΩ·cm^2^, unit axoplasm resistance *R*_i_ – within 79… 314 Ω·cm, and unit membrane capacity *C_m_* = 1 *μ*F/cm^2^ is nearly uniform between all species. *R_m_* determines how quickly the current spreading through a neurite attenuates due to leakage through its membrane. In *C. elegans* the value of *R_m_* is quite high in comparison to other species (typically 1-50 kΩ·cm^2^), which favors farther signal propagation. Cell membrane resting potential, remaining nearly constant at rest and changing in response to incoming signals, is also of high importance. Electrophysiologic data about resting potential of *C. elegans* neurons is available for only a few of them. Registered values are settled within the interval from −20 to −50 mV ^31^. For a larger nematode *A. suum* measurement of the resting potential is easier. It is known to be about −40 mV. We decided to use this value since it is within the interval of values observed for *C. elegans.* The mechanism of passive signal propagation and its attenuation due to outflow through leakage channels (defined by the value of *R_m_*) is built into NEURON. The data which we have described in this section were used for the construction of the model of *C. elegans* neuron.

Every compartment of a neuron model in NEURON simulator is represented by a conical frustum defined by (*x_i_*, *y_i_*, *z_i_*, *d_i_*) and (*x*_*i*+1_, *y*_*i*+1_, *z*_*i*+1_, *d*_*i*+1_), where *x, y, z* are 3D coordinates of the center of each base circle and d is its diameter. Every compartment may possess its own electrochemical and morphological parameters, ion channels and other mechanisms, and is usually connected with previous and next compartment, or compartments – in case of branching. Any two points of two different neurons can be connected by a synapse or a gap junction. Usually a synapse conducts signal only in one direction – from pre-synaptic neuron to a post-synaptic one, and a gap junction – in both ways. A typical time required for a signal to pass through a synapse is about 2 ms and for a gap junction – 0.2 *ms.* Both mechanisms are available in NEURON.

### 2.2. On electrophysiology of C. elegans muscles

#### 2.2.1. Ion channels

In contrast to *C. elegans* neurons, at least two major classes of its muscles (pharyngeal and body wall muscles) are capable of firing APs, driven by Ca^2+^ entry through the EGL-19 L-type Ca^2+^ channel with contribution from the CCA-1 T-type Ca^2+^ channel. The latter is activated by small depolarizations from resting potential, lifts membrane potential until EGL-19 can be activated and inactivates quickly^17^. The duration of APs in a body wall muscle and in a pharyngeal muscle is about 20 ms and 150 ms correspondingly^7,11^, which is much longer than in vertebrate skeletal muscle (<2 ms). T-type and L-type Ca^2+^ channels are responsible for depolarization and long plateau, correspondingly. The plateau is interrupted by a rapid repolarization depending on the function of voltage-gated K+ channels, which differ in case of pharyngeal and body wall muscles. Here we focus on pharyngeal muscles, in which the role of key voltage-gated K+ channel belongs to EXP-2^7^. L-type EGL-19 Ca^2+^ and Kv-type EXP-2 channels are subject to this research and are considered in the ”Results” section. The similar approach can be applied to body wall muscles as well.

The following parameters responsible for equilibrium potentials and ion channel conductances ion channel were used:

|  |  |
| --- | --- |
| *gmax_KvEXP2* | – maximal EXP-2 conductance, [$S/cm^2$] |
| *gmax_CaL* | – maximal EGL-19 conductance, [$S/cm^2$] |
| *gmax_pas* | – maximal conductance of the passive (leak) current, [$S/cm^2$] |
| *ek* | – potassium equilibrium potential, [$mV$] |
| *eca* | – calcium equilibrium potential, [$mV$] |
| *e_pas* | – equilibrium potential of the passive current, [$mV$] |

A recording of the membrane potential of an active pharyngeal muscle cell was performed in the work of M.W. Davis and colleagues (at Fig.5a)^12^. We used their data to optimize the difference between a real and simulated AP and compare these curves (in the ‘‘Results” section). According to C. ELEGANS II book^13^, this shape of AP is very similar to those recorded from *Ascaris suum* pharyngeal muscle cell^14^ and resemble APs of vertebrate heart muscle cells^15^.

#### 2.2.2. Pharyngeal muscle cell membrane resting potential

Modeling of a cell electrophysiologic activity requires knowing the value of its membrane resting potential. According to the work of C.M. Rogers with colleagues^16^, it was −69±1 mV. K.A. Steger and colleagues obtained the value about −73 mV^17^. Other studies^12,18^ reported the values from −40 to −50 mV. This difference probably might be due to the recording from different muscle cells of a pharynx or using different recording techniques and/or equipment. We decided to use −50 *mV* since this value is observed at action potential recordings^12^ performed with high time resolution which we use to minimize the difference between the real and simulated action potential shapes.

#### 2.2.3. Leak conductance and equilibrium potential

Passive leak current also makes and impact on the shape of the AP and depends on pharyngeal muscle cell leak conductance and equilibrium potential for the leakage current. These values were reported by B. Shtonda and L. Avery^7^: 22.8±5.2 MΩ and −53.6±4.1 mV correspondingly. Pharynx capacitance is also known, 362.6±36.8 pF, which allows one to estimate unit leak conductance (in S/cm^2^), required for modeling in NEURON. It is well-known that unit membrane capacitance for the majority of neurons, muscle and other types of cells is about 1 μF/cm^2^. The area of a pharynx surface can be estimated as (362 pF)/(1 *μ*F/cm^2^) = 0.363· 10^−3^ cm^2^. Finally, the unit leak conductance = (1/22.8 MΩ)/(0.363· 10^−3^ cm^2^) = 1.24·10^−4^ S/cm^2^. A leak conductance of body muscle cells, mentioned in the work of J.H. Boyle^19^, is equal to −22 S/F, which is equivalent to 2.2·10^−5^ S/cm^2^ (or 45 kΩ·cm^2^).

## 3. Results and Discussion

### 3.1. Basic neuron and signal conductance

We have created a model of *C. elegans* neuron using the NEURON simulation environment using the following parameters: soma diameter – 2 μm, neurite diameter – 0.2 μm, neurite length – 1 mm (see Fig. 1).

**Fig. 1.**
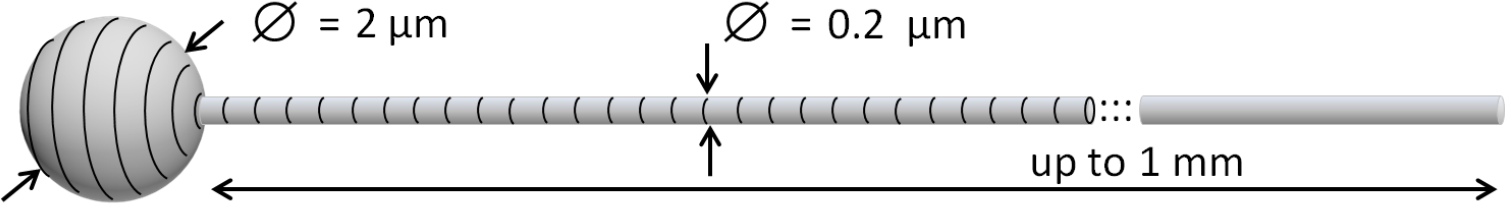
3D view of *C. elegans* neuron and neurite representation in NEURON.

The resting potential was equal to −40 mV, specific membrane resistance and unit axoplasm resistance were varied within the limits mentioned in the section 2.1. For a variety of considered combinations of parameters the signal propagation length constant λ and velocity of electrical excitation propagation θ were estimated using the following formulas:

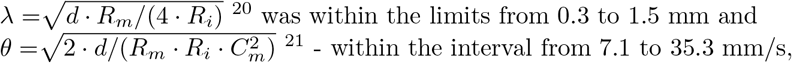

where *d* is a neurite diameter.

The experimental data on propagation of a signal along 350 *μ*m long *C. elegans* neurite are available from work of Suzuki and colleagues^22^. Using them we estimated the velocity of this process^10^. Obtained interval of values, 10… 25 mm/s, is in a good accordance with our initial estimate for *θ*.

To verify this in practice we simulated an injection of a current into one end of the neurite and detected the response at another. The current causing the signal in the work^22^ was generated by mechanosensory MEC-4 Na+ channel. In our simulation we took into account that the ionic current through the single channel was equal to 1.6 pA and 14… 25 channels were simultaneously open^23^ (during nearly 10 ms). The signal registered at another end of neurite, after passing a distance of 1 mm, came with a latency of nearly 0.1 s (which corresponds to a 10 mm/s velocity) and with a peak value of the membrane potential equal to −28 mV (half of the peak value was reached in 0.04 s). In some cases this is enough for activation of a post-synaptic cell. The proposed model reproduces the basic features of a neuron quite well and might be used further as an initial point for development of *C. elegans* neural network.

### 3.2. Modeling of a pharyngeal muscle cell

Among two largest pharyngeal muscle cells – ”pm3” and ”pm5”, we have chosen the first one for modeling, because it contracts in an ordinary way, all-at-once, whereas ”pm5” does it via anterior to posterior peristalsis wave. Actually ”pm3” is the common name for three cells (dorsal, L-subventral and R-subventral) composing a muscular tube cylinder, along with three smaller ”mc1” marginal cells^g^. According to the microphotograph, this cylinder is about 43 μm long and 15 μm in diameter. We approximate a single ”pm3” cell (any of three) with a cylinder of the same length and 7.7 *μ*m in diameter (see Fig.2).

**Fig. 2.**
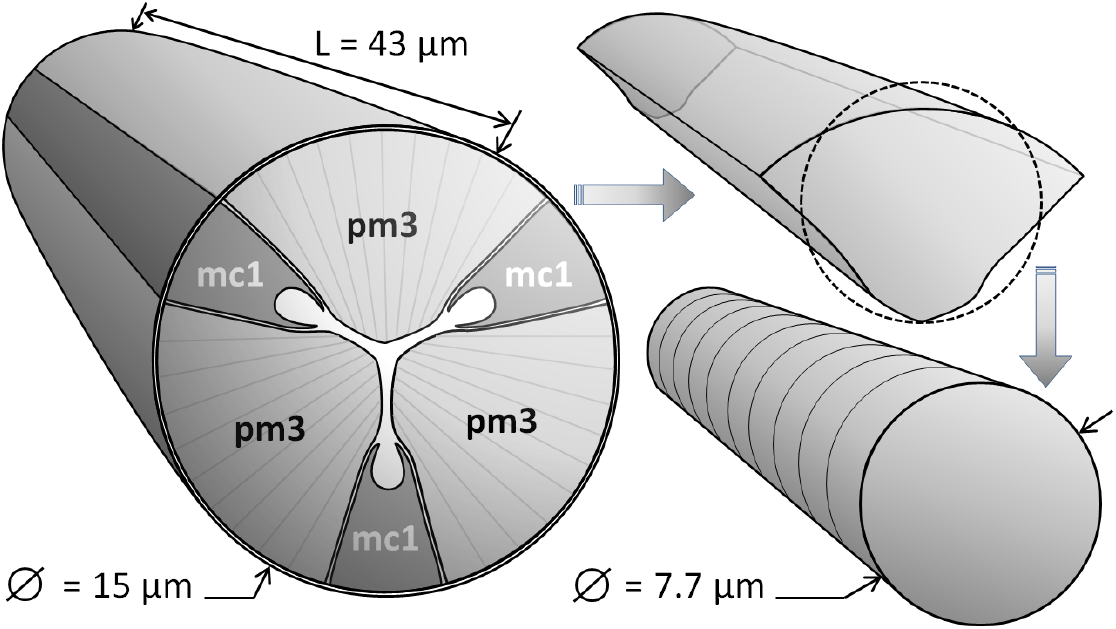
A simplified 3D view of ”pm3” and ”mc1” cells (left), of one of three pm3 cells separately (upper right) and of its representation in NEURON (lower right).

After definition of the ”pm3” pharyngeal muscle cell geometry we added and tuned EXP-2 and EGL-19 ion channels models (separately) and finally optimized the model which includes both channels simultaneously.

#### 3.2.1. L-type EGL-19 Ca^2+^ channel

Among existing models of Ca^2+^ channels the most related one was developed for a cardiac muscle. It is a smooth non-striated muscle as well as *C. elegans* pharyngeal muscles (however, its body wall muscles are striated)^h^. We reproduced the corresponding Markov model based on the state diagram, differential equations and parameters from the work of V.E. Bondarenko and colleagues^24^. It was implemented as a MOD-file (written in Neuron Model Description Language), which was compiled to become accessible within NEURON simulation environment. To verify the model, we reproduced *in silico* the voltage-clamp experiments equivalent to those presented in Fig.3A,B of the same work^24^. Our calculations resulted in the similar dependence of Ca^2+^ peak current on the voltage, including the position of maximal magnitude of the current corresponding to 0 mV. (see Fig.3A).

**Fig. 3.**
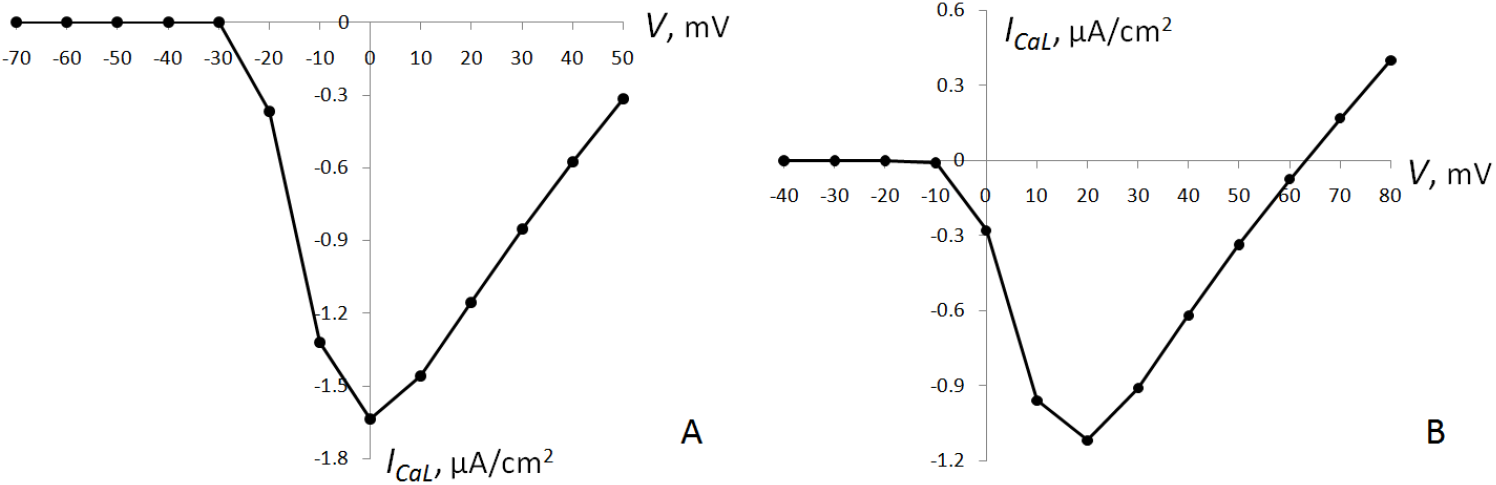
Voltage-current curves for the original (A) and modified (B) versions of Ca^2+^ L-type ion channel Markov model.

However, EGL-19 Ca^2+^ L-type channel in *C. elegans,* although it has a similar shape of current-voltage dependence, has its maximal magnitude of current located at nearly 20 mV, not 0 (see Fig.3D (”wild-type” curve) in the work of V. Laine and colleagues^25^). Changing the parameters of L-type calcium channel, one by one, we have found that there is a simple modification that moves the I-V dependence 20 mV to the right so that it becomes similar to the real current-voltage relationship for EGL-19 (see Fig.3B.). The changes which thus convert the model of cardiac myocyte’s L-type Ca^2+^ channel into the model of EGL-19 L-type Ca^2+^ channel are the following: formulae (32) and (33) for *α* and *β* from the work^24^ should be modified according to this schema:

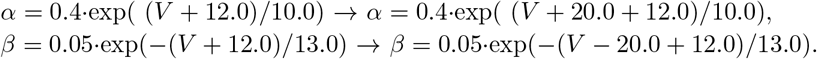

#### 3.2.2. Kv-type EXP-2 channel

EXP-2 voltage-gated K^2+^ channel has no existing Markov model. However, its most closely related ion channel belongs to the Kv-type family (Kv 11.1, human HERG K^2+^ channel), with a number of models proposed^26^. HERG channel is functionally distinct and very similar to EXP-2 – both activate slowly and inactivate very rapidly in response to depolarization. Although they are structurally unrelated, EXP-2 kinetic properties (which are the key to Markov model) make it an inward rectifier that resembles HERG channel^27^. Although they are similar, but not equivalent. For example, EXP-2 single-channel conductance is 5- to 10-fold larger than that of HERG and most Kv-type K+ channels^27^, so we decided to optimize a set of parameters used in the Markov model of HERG channel to reconfigure it into EXP-2. Here is the list of parameters defining transitions between states of the Markov model (one open, tree closed and one inactivated) from the work of R. Mazhari and colleagues^28^, in its original form (left) and with our modifications (right), introducing 16 parameters (*p*_1_-*p*_1_β) which will be optimized (V is the membrane voltage):

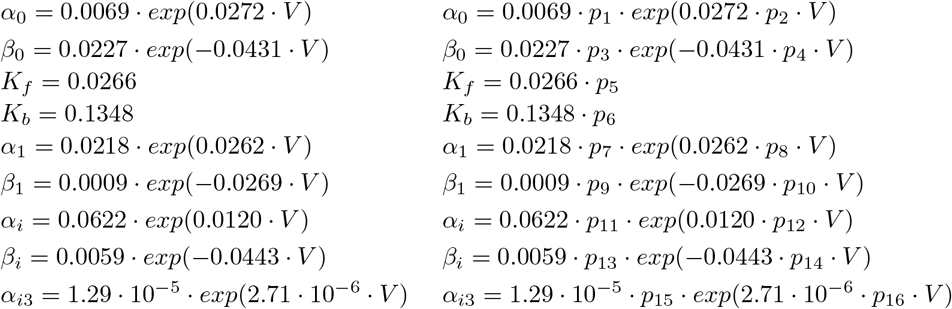

Electophysiologic properties of EXP-2 were previously studied in the experiments with expression of these channels in *Xenopus* oocytes^27^. Fig.3A of the corresponding paper presents a series of currents recorded from whole oocytes via two-electrode voltage clamp technique. In each recording EXP-2 channels were inactivated by 1 s prepulses to 20 mV, then allowed to recover from inactivation at −80 mV for 40 ms and finally were affected by 1 s pulses to voltages ranging from – 120 to 60 mV (with 10 mV increments)^27^. We digitized three curves (corresponding to −120 mV, −80 mV and +60 mV pulses) to use them for optimization of EXP-2 parameters via an equivalent *in silico* experiment.

Our optimization algorithm implemented in HOC (NEURON’s ‘native’ language) performs stochastic directed search over the space of parameters. At every step of the main optimization loop, small changes are applied to the parameters each time randomly chosen their full set. The current through the voltage clamp is calculated as a result of simulation and then compared with a curve based on experimental data. The modulus of difference between each pair of points corresponding to the same moment of time is calculated and accumulated over the whole time interval (total difference). The new combination of parameters is either accepted or declined, with a probability depending on the relation between the previous and the new values of the total differences. The process continues until the convergence or lasts for a fixed number of iterations, usually about 100000, which may take up to several hours on a modern desktop PC. Previously we successfully used the similar algorithm for optimization of parameters describing a kinetic scheme of transitions between protein states^29^.

Optimization experiments showed that all three curves (corresponding to voltage steps to −120, −80 and +60 mV), being optimized separately, were reproduced nearly perfectly (with less than 2.8% difference). However, when we used this or that set of optimized parameters in the simulation with another value of voltage step, the dependence of current on time became rather satisfactory than good. To solve this we optimized two curves (corresponding to voltage steps to −120 and −80 mV) simultaneously, minimizing the total difference between each real and simulated curve. In this case we have got significantly better results, with a difference about 11.4%. The obtained set of parameters was used to generate a series of curves (shown in Fig.4), which reproduce the corresponding experiment quite well. We also calculated the I-V relationship via *in silico* reproduction of the experiment presented in Fig.3C (”100 mM” curve) from the work of Fleischhauer and colleagues^27^. The shape of I-V curve obtained in our simulation resembles the real one (see Fig. 5).

**Fig. 4.**
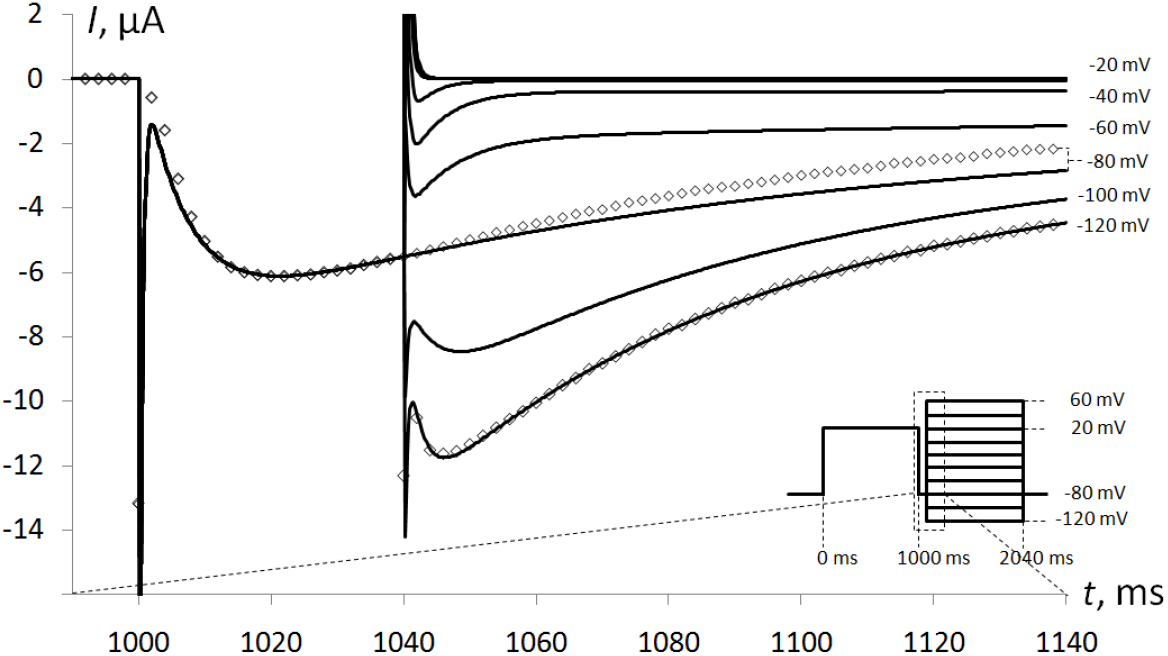
A set of curves displaying currents through patch-clamp corresponding to steps to voltages ranging from −120 to +60 mV with 20 mV increments – *in silico* reproduction of the experiment described in the work of Fleischhauer and colleagues, at their Fig.3A. Solid lines display the results of simulation, diamonds – digitized points of −120 and −80 curves obtained from the experiment.

**Fig. 5.**
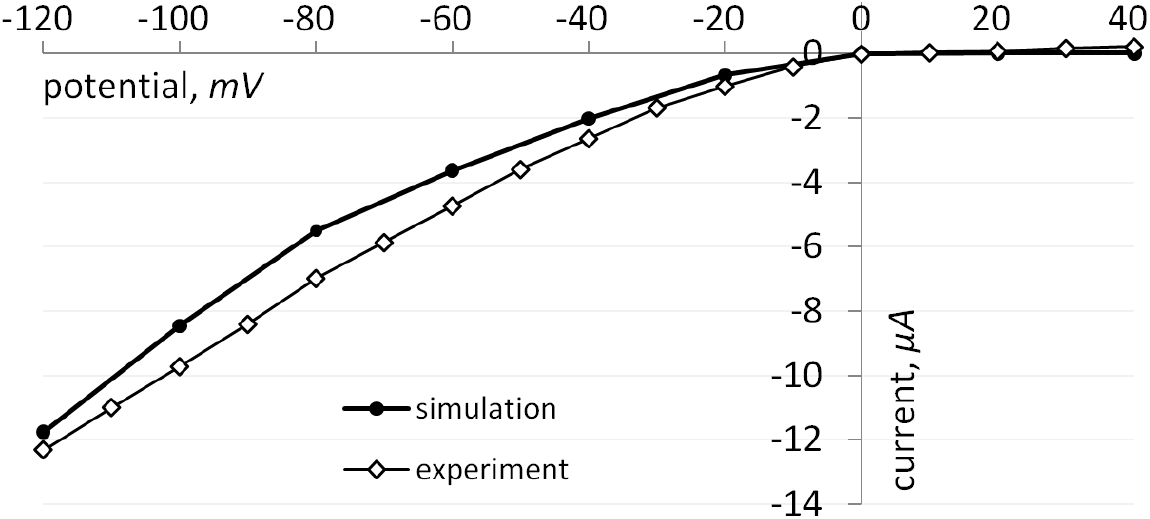
I-V relationship for EXP-2 at 100 mM extracellular KCl concentration, obtained in the experiment by Fleischhauer and colleagues and in our simulation.

All our simulations based on oocytes were performed with the value of *e_k_* = -2 mV. It was calculated via the Nernst equation, taking T = 20 C, extracellular K+ concentration = 100 mM^27^ and intracellular K+ concentration = 108.6 mM (see Tables 2 and 3 of the paper^32^). The resulting parameters which thus convert the model of HERG channel to the model of EXP-2 are listed in Table 1 (column ‘Simulation 1”).

**Table 1.**
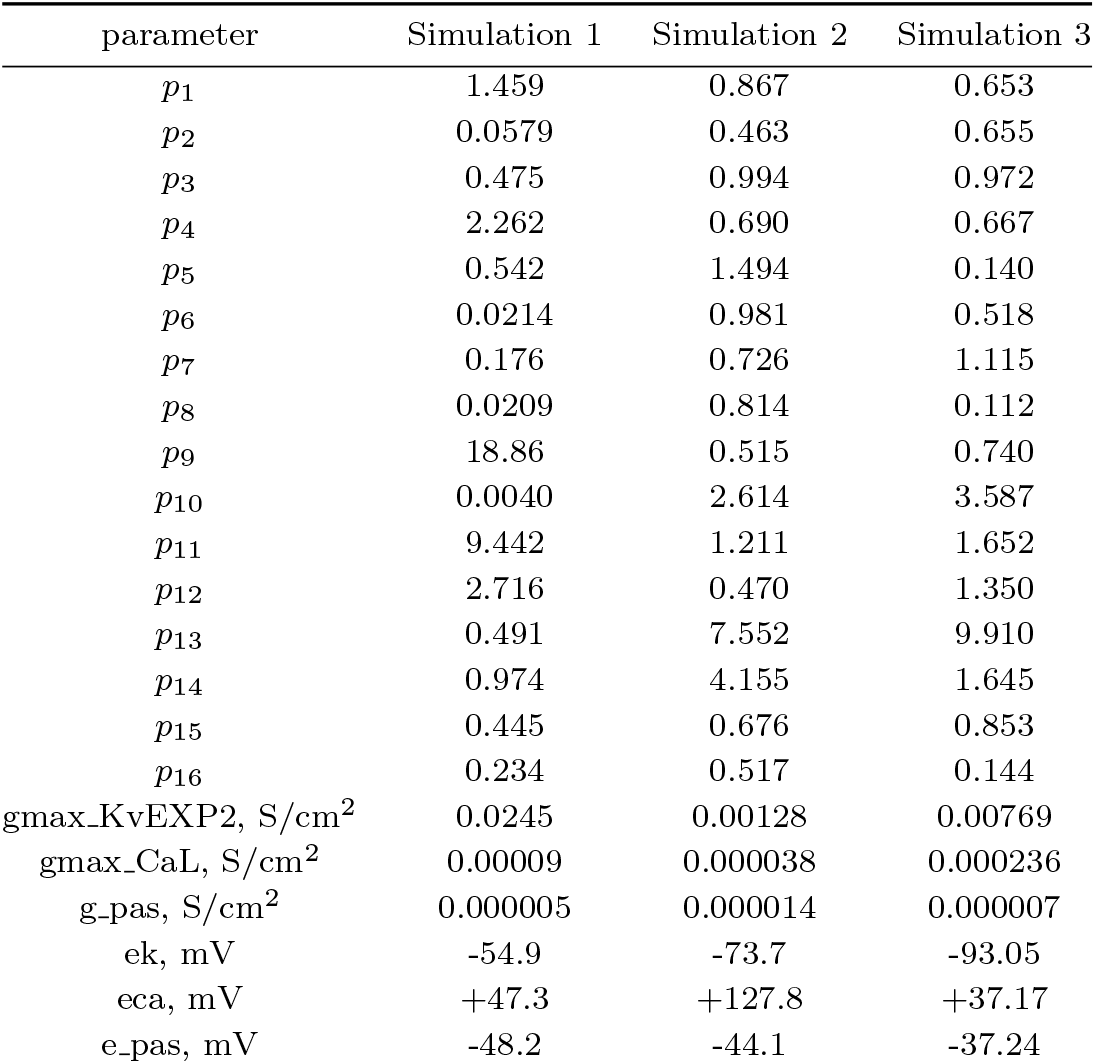
Parameters obtained as a result of optimization.

| parameter | Simulation 1 | Simulation 2 | Simulation 3 |
| --- | --- | --- | --- |
| $p_1$ | 1.459 | 0.867 | 0.653 |
| $p_2$ | 0.0579 | 0.463 | 0.655 |
| $p_3$ | 0.475 | 0.994 | 0.972 |
| $p_4$ | 2.262 | 0.690 | 0.667 |
| $p_5$ | 0.542 | 1.494 | 0.140 |
| $p_6$ | 0.0214 | 0.981 | 0.518 |
| $p_7$ | 0.176 | 0.726 | 1.115 |
| $p_8$ | 0.0209 | 0.814 | 0.112 |
| $p_9$ | 18.86 | 0.515 | 0.740 |
| $p_{10}$ | 0.0040 | 2.614 | 3.587 |
| $p_{11}$ | 9.442 | 1.211 | 1.652 |
| $p_{12}$ | 2.716 | 0.470 | 1.350 |
| $p_{13}$ | 0.491 | 7.552 | 9.910 |
| $p_{14}$ | 0.974 | 4.155 | 1.645 |
| $p_{15}$ | 0.445 | 0.676 | 0.853 |
| $p_{16}$ | 0.234 | 0.517 | 0.144 |
| $g_{\max\_KvEXP2}$, S/cm<sup>2</sup> | 0.0245 | 0.00128 | 0.00769 |
| $g_{\max\_CaL}$, S/cm<sup>2</sup> | 0.00009 | 0.000038 | 0.000236 |
| $g_{\_pas}$, S/cm<sup>2</sup> | 0.000005 | 0.000014 | 0.000007 |
| $e_k$, mV | -54.9 | -73.7 | -93.05 |
| $e_{ca}$, mV | +47.3 | +127.8 | +37.17 |
| $e_{\_pas}$, mV | -48.2 | -44.1 | -37.24 |

#### 3.2.3. The final model provided with EGL-19 and EXP-2 ion channels

When EGL-19 and EXP-2 channels were ready, additional optimization of passive, calcium and sodium equilibrium potentials and conductances was performed. Minimization of the difference between the real and simulated pharyngeal muscle’s AP curves resulted in the shape which was relatively close to that presented in Fig.7 of the work of Shtonda and Avery^7^. However, it didn’t reproduced the hyperpolarization phase following the rapid repolarization after the end of the plateau (see Fig.6A, ‘Simulation 1”).

**Fig. 6.**
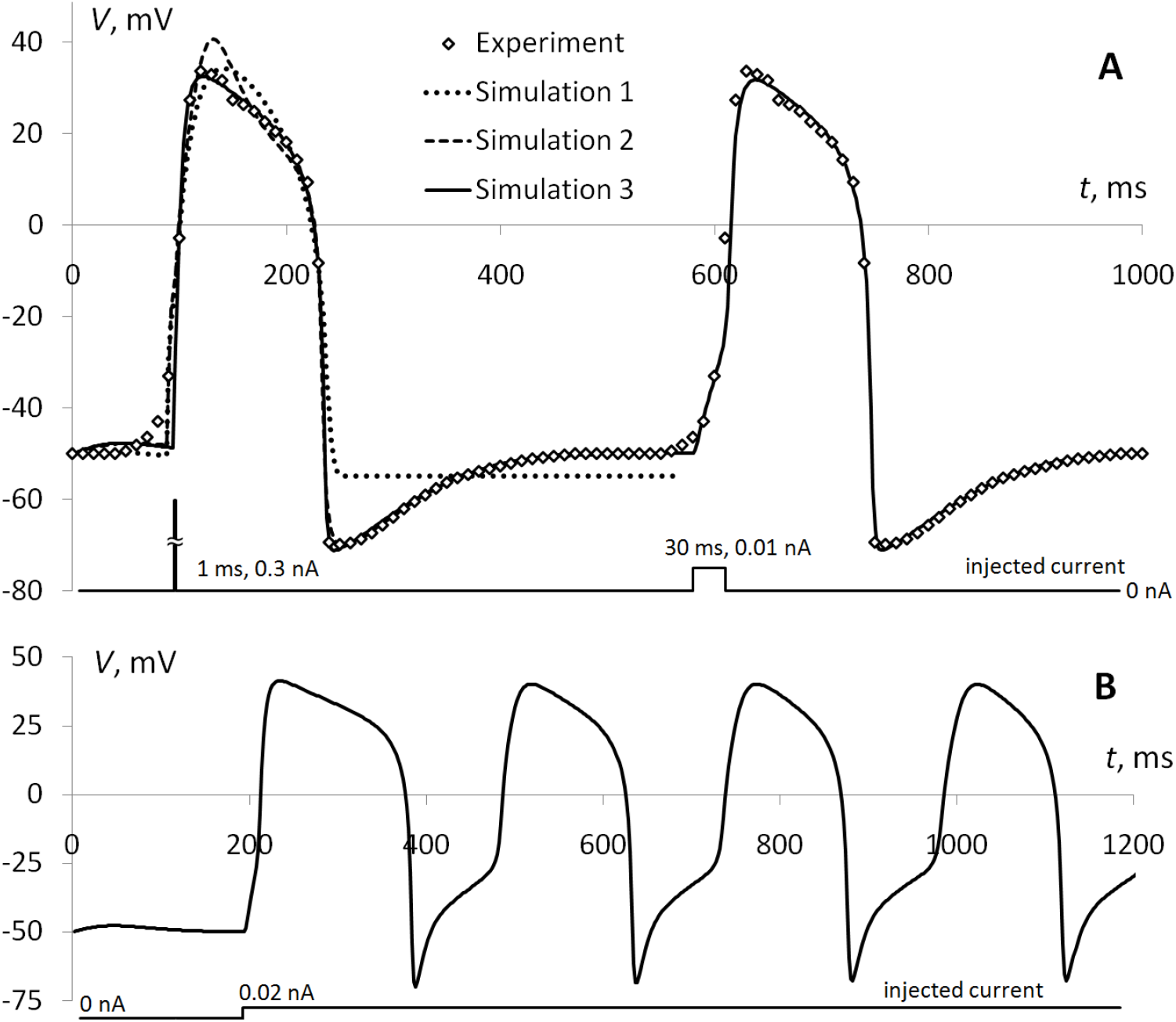
A. The curve ‘‘Experiment” displays pharyngeal action potential recorded in experiment performed by M.W. Davis and colleagues, mentioned earlier (data was digitized from their Fig.6a). ‘‘Simulation 1” shows the shape of AP obtained as a result of optimization without changing of *p*1-*p*16 parameters (tuned for EXP-2 channel). ‘‘Simulation 2” and ‘‘Simulation 3” display two different variants of APs, obtained with *p*1-*p*16 included into optimization. **B**. The train of pharyngeal APs induced by a continious injection of 0.02 nA current.

Finally we performed the optimization in which the parameters p_1_-p_1_6 were additionally included. Convergence took a long time – even after 500000 iterations small further improvements were observed. However, the resulting shape of the simulated AP fits the real one almost perfectly, with a difference only in the phase of initiation. The latter corresponds to T-type CCA-1 Ca^2+^ channel, which is not yet implemented in our model (see Fig.6A, first impulse). Interestingly, changing the profile of injected current from 0.3 nA during 1 ms to 0.01 nA during 30 ms (see Fig.6A, ‘Simulation 3”, second impulse) makes the shape of the simulated AP similar to the shape of the real one, even at the initial stage. Since we inject the current artificially, imitating the action of CCA-1 channel of a real muscular cell, noticeably better similarity of AP in the second case might speak well for the shape of the real current flowing through this channel.

The difference between the shape of a real AP and ‘‘Simulation 3”, the best one, is about 3.2%. Results obtained in several optimization runs showed that equilibrium potentials for K+ and passive leakage currents are usually within a narrow interval, whereas the one for Ca^2+^ is much wider – a spectrum of solutions is available. It other words, optimization with a fixed value of a calcium resting potential will probably still lead to a good solution.

Further studies of the model revealed additional resemblance between the simulated and real ”pm3” cell. It is known that usually the rate of pharyngeal pumping varies from about 0.7 Hz to 4.4 Hz, depending on the presence of food and other conditions. Each action potential is initiated by the incoming signal and corresponds to a single pump. We have simulated the injection of impulses of current (0.02 nA, 30 ms) increasing their frequency and finally the sequence of pulses became a single continuous current (Fig.6B). The maximal frequency of APs observed in these simulations was equal to 4.3 Hz and didn’t grew anymore, even with the increase of the amplitude of the current. With this we have got a very close value of the pharyngeal pumping rate limit and obtained the curve which is known as the train of pharyngeal APs, with the first peak longer than all subsequent, which is presented and described in the Fig.3a from the work^i^.

## 4. Conclusion

In the present work we proposed the computational models of a typical *C. elegans* neuron and of a pharyngeal muscle, based on the basis of the available experimental data, including elecrophysiological properties and key ion channels. Obtained models are in accordance with known facts about simulated objects and reproduce their properties really well. We initiated this work since, to our knowledge, at that moment there were no other *C.* elegans-related models of this kind taking into account the specificity of the considered cell types based on reproduction of their deep inner mechanisms like ion channels.

We position the proposed models as a basic ‘building blocks” for both the development of more sophisticated models of individual cells and for construction of networks composed of these elements, reproducing the architecture of real system. Implementation of all developed source codes in terms of the NEURON’s programming language allows them to be available to a wide audience since it is used all over the world for scientific and educational purposes.

Realistic modeling of a complex system’s basic building blocks is usually performed to make high-quality predictions on its larger-scale properties or to simulate its behavior. Using the model of a pharyngeal muscle cell we revealed that it has the upper limit of action potentials frequency matching the value known from the experiment. And for the case of a neuron the transmission of a signal along 1 mm long neurite was simulated in accordance with all key electrophysiologic parameters known for this system.

However, our expectations on these models are significanlty larger and hopefully will allow to unveil their full potential in near future. Our recently developed simulation engine called Sibernetic^30j^ performs realistic 3D simulation of worm body crawling^k^ and swimming^l^. Using the similar techniques, we plan to supplement it with a physical model of pharyngeal muscles supplemented with their electrophysiological model and driven by simulated electrical activity of corresponding neurons, based on the models proposed in the present work.

## Acknowledgments

The work of A. Palyanov and K. Samoilova was supported by Russian Federation President grant MK-5714.2015.9.

## Footnotes

a https://www.humanbrainproject.eu

b http://www.humanconnectomeproject.org

c http://www.openworm.org

d http://www.wormatlas.org/hermaphrodite/pharynx/mainframe.htm

e http://neuron.yale.edu/neuron

f http://genesis-sim.org

g http://www.wormatlas.org/hermaphrodite/pharynx/mainframe.htm, PhaFIG 6 and 7A

h http://www.wormatlas.org/hermaphrodite/muscleintro/mainframe.htm

i http://www.wormbook.org/chapters/www_feeding/feeding.html

j http://sibernetic.org

k https://www.youtube.com/watch?v=JwG5PfDIoU

l https://www.youtube.com/watch?v=AQAme81B4K0

---

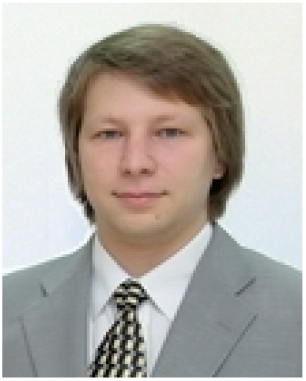
**Andrey Palyanov** received his MSc degree in biological and chemical physics from Novosibirsk State University, Russia, in 2004 and Ph.D. degree in chemical physics from Voevodsky Institute of Chemical Kinetics and Combustion in 2008. During his Ph.D. studies he worked in Kutateladze Institute of Thermophysics, Laboratory of Simulation, and in Universite Louis Pasteur, Laboratory of Biophysical Chemistry (headed by Martin Karplus). From 2008 to present time he works in Ershov Institute of Informatics Systems, Laboratory of Complex Systems Simulation as a senior researcher. In 2011 he joined the international OpenWorm Project aimed at creating a virtual *C. elegans* nematode in a computer, and belongs to core team members of the project. And since 2015 he also collaborates with Novosibirsk State University, Laboratory of Structural Bioinformatics and Molecular Modeling.

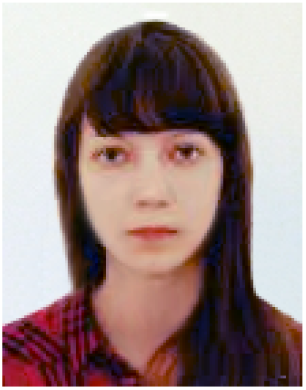
**Khristina Samoilova** received her Bachelor’s degree in Mathematics and Computer Science from Novosibirsk State University, Russia, in 2015. During her scientific practice she worked in A.P. Ershov Institute of Informatics Systems in the Laboratory of Complex Systems Simulations, in the field of application of computational methods to the problem of *C. elegans* nervous system investigation. In 2017 she has obtained her Master’s degree in Inverse and Ill-posed Problems from the same university and continues her scientific work.

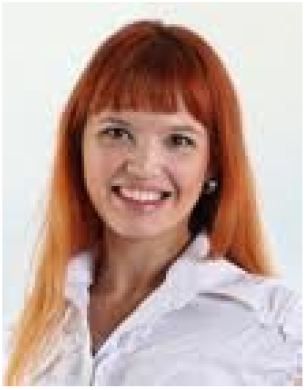
**Natalia Palyanova** received her Specialist’s degree in Physiology (MSc/MA qualification in Biology) from Novosibirsk State University (Department of Natural Sciences), Russia, in 2006. She works in the Institute of Molecular Biology and Biophysics as a junior researcher in the Laboratory of Central Mechanisms of Regulation and Control. Currently she finishes her Ph.D. thesis in the field of neuroprotection against oxygen-glucose deprivation. She participates in this project as an expert in neurobiology.

